# Vascular cognitive impairment in the mouse reshapes visual, spatial network functional connectivity

**DOI:** 10.1101/2020.11.04.366294

**Authors:** Gerard R Hall, Philipp Boehm-Sturm, Ulrich Dirnagl, Carsten Finke, Marco Foddis, Christoph Harms, Stefan Paul Koch, Joseph Kuchling, Christopher R Madan, Susanne Mueller, Celeste Sassi, Stamatios N Sotiropoulos, Rebecca C Trueman, Marcus Wallis, Ferah Yildirim, Tracy D Farr

## Abstract

Connectome analysis of neuroimaging data is a rapidly expanding field to identify disease specific biomarkers. Structural diffusion MRI connectivity has been useful in individuals with radiological features of small vessel disease, such as white matter hyperintensities. Global efficiency, a network metric calculated from the structural connectome, is an excellent predictor of cognitive decline. To dissect the biological underpinning of these changes, animal models are required. We tested whether the structural connectome is altered in a mouse model of vascular cognitive impairment. White matter damage was more pronounced by 6 compared to 3 months. Global efficiency remained intact, but the visual association cortex exhibited increased structural connectivity with other brain regions. Exploratory resting state functional MRI connectivity analysis revealed diminished default mode network activity in the model compared to shams. Further perturbations were observed in a primarily cortical hub and the retrosplenial and visual cortices, and the hippocampus were the most affected nodes. Behavioural deficits were observed in the cued water maze, supporting the suggestion that the visual and spatial memory networks are affected. We demonstrate specific circuitry is rendered vulnerable to vascular stress in the mouse, and the model will be useful to examine pathophysiological mechanisms of small vessel disease.

**Graphical abstract:** 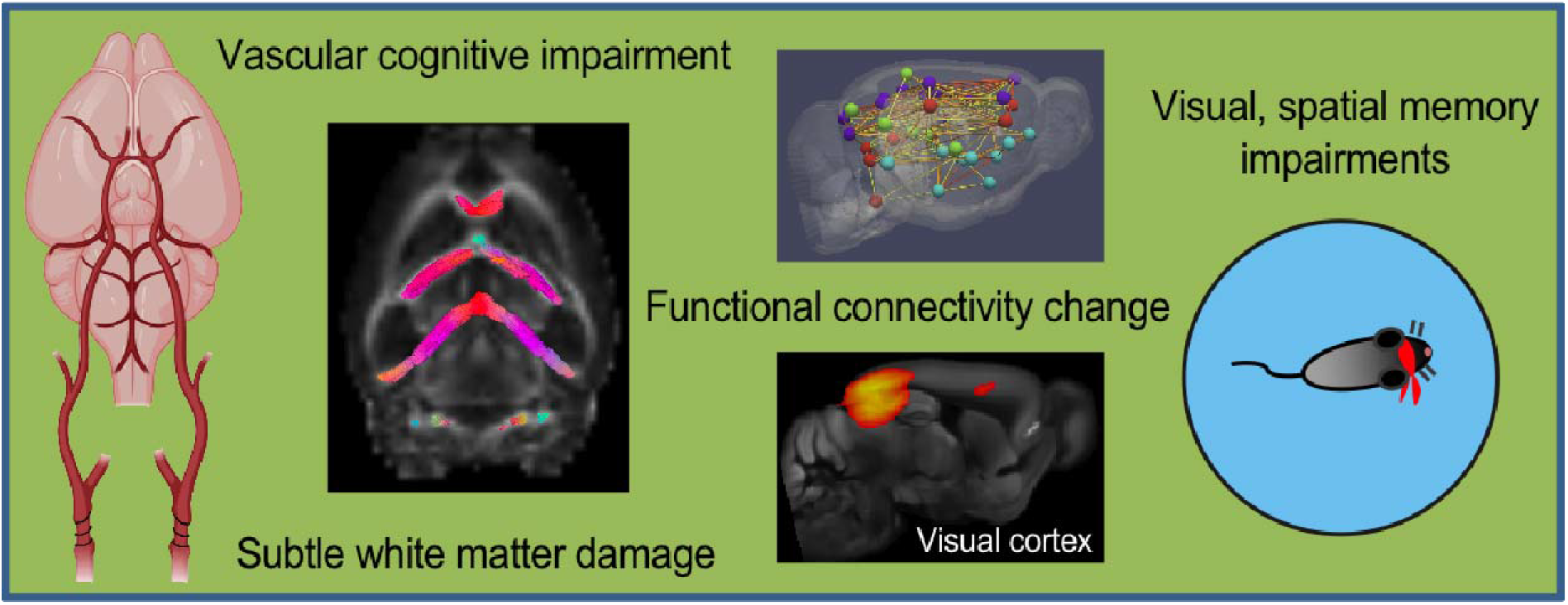

## Introduction

Neuroimaging is an important tool to non-invasively characterize the brain and examination of whole brain structural and functional connections is a rapidly evolving field. Connectomics aims to improve our understanding of healthy brain architecture and offers the potential to improve diagnosis, patient stratification, and outcome prediction in disease conditions. There is a growing body of evidence that suggests connectivity patterns may be disease specific, reviewed in (Hallett et al., 2020; Lazarou et al., 2019). Structural connectivity analysis from diffusion MRI (dMRI) has been particularly useful as a biomarker for small vessel disease due to the prevalence of white matter changes in these individuals (Qin et al., 2020; Reijmer et al., 2016). More specifically, reduced global efficiency (a quantitative overall network metric that characterizes a feature of the structural connectome) has been consistently reported as a good predictor of cognitive decline and conversion to dementia in patients with radiological features of small vessel disease (Lawrence et al., 2018; ter Telgte et al., 2018; Tuladhar et al., 2016). These structural network disruptions may be accompanied by functional connectivity alterations that can be measured using resting state functional MRI (rsfMRI). Indeed, decreases in functional connectivity have been reported in frontoparietal (Schaefer et al., 2014) and frontotemporal regions (Sun et al., 2014; Yu et al., 2015) in individuals with small vessel disease. The default mode network (DMN) refers to a collection of structures (posterior cingulate, precuneus, medial prefrontal and inferior parietal cortices) with highly correlated activity patterns during internally directed tasks; it is a fundamental brain network that appears to be disrupted in a variety of neurological diseases, including those with vascular origins (Papma et al., 2013).

Animal models can bridge the gap between macroscale imaging-based connectivity analysis, and the microscale cellular and molecular processes that underpin it. Recent work has linked mouse functional connectivity patterns to distinct axonal projections (Grandjean et al., 2017; Whitesell et al., 2020) and some groups have reported gene specific connectivity alterations (Mechling et al., 2016; Pagani et al., 2019). Though functional connectivity analysis of rsfMRI is lagging in rodents compared to humans, mouse DMN connectivity appears to decline with age (Egimendia et al., 2019). Reports are also emerging in mouse models of Alzheimer’s disease that suggest reduced functional connectivity in the cortex precedes amyloid plaque deposition (Grandjean et al., 2014), and is correlated with tau aggregation (Green et al., 2019). The purpose of the present study was to use a hypothesis free approach to examine functional and structural connectivity change, including global efficiency, in a mouse model of vascular cognitive impairment that is characterized by white matter damage and spatial memory impairments, reviewed in (Duncombe et al., 2017). We show substantial functional connectivity alterations in cortical hubs, particularly those associated with the visual and spatial memory system. This was accompanied by predictable behavioural deficits with little evidence of structural connectivity breakdown.

## Materials and Methods

### Animals, surgical procedures and experimental design

Experiments were performed according to the Landesamt für Gesundheit und Soziales and conformed to ARRIVE guidelines. C57/BL6J mice (8 weeks of age, Charles River, Germany) were housed in groups of 6 in filter top cages with tubes, nesting, and chew blocks under a 12 hr light/dark cycle (temperature: 22 ± 2 °C, humidity: 55 ± 10 %). They had *ad libitum* access to food and water. Animals were anaesthetized with isoflurane (70:30 nitrous oxide:oxygen) and body temperature was maintained at 37 ± 0.2 °C during all procedures. The hypoperfusion or sham procedure involved wrapping a non-magnetic microcoil (160 or 500 μm, hypoperfused (n=11) or sham (n=10), respectively) around a carotid artery. The procedure was repeated for the other carotid artery the following day. Pain relief was provided in the drinking water (6 mg/mL of Paracetamol) 3 days prior to surgery and for up to a week following. There was high mortality in the hypoperfused group: 2 animals died within 48 hrs, and 3 others at 9, 28, and 32 days post-surgery. MRI was conducted at 24 hrs (T_2_, CBF), and 3 (T_2_, CBF, dMRI, MRA, spectroscopy) and 6 months post-surgery (T_2_, CBF, dMRI, MRA, spectroscopy, rsfMRI). Novel object recognition and the Morris water maze cued and place tasks were performed at 6 months. Analysis of all data was performed blind.

### Behavioural testing

All behavioural tests were recorded and analysed with a computerized automated tracking system (VideoMot 2, TSE Systems, Germany). For novel object recognition, mice were habituated to an empty cage for 10 min on the day prior to testing. During the first trial (10 min) mice were positioned facing away from two identical wooden objects (kept in bedding from unfamiliar mice) secured to the floor. Two hours later, the right object was replaced (stored in bedding from a different group of unfamiliar mice) and mice were allowed to explore for 5 min. Total time spent with each object was calculated, and data was presented as discrimination ratio (right object / (right object + left object)).

The Morris water maze (120 cm diameter circular pool) contained visual cues. A cue task was performed for 4 days with a visible platform in the same location (3 trials of maximum 90 s). If mice failed to escape, they were guided to the platform. Subsequently, a place task was used for 7 days in which the escape platform was submerged (4 trials, 20 min apart, with randomization of entry point). Animals were collected 10 s after locating the platform. A probe trial without a platform was performed on the 7^th^ and 8^th^ day. Escape latency (s), total distance travelled (cm), swim speed (cm/s), and total time spent in each quadrant (s) were measured, and daily data was pooled.

### MRI measurements

All MRI measurements were performed using a 7T system (Bruker BioSpin, Germany) under isoflurane anaesthesia (including for the rsfMRI, but at low levels) and temperature and respiration were monitored (SA Instruments, USA). Cerebral blood flow (CBF) was measured with a 72 mm diameter transmit volume coil (RAPID Biomedical, Germany) and mouse quadrature receive surface coil (Bruker BioSpin) using a 2D Flow-Sensitive Alternating Inversion Recovery (FAIR) sequence (repetition time/echo time (TR/TE): 12 000/35.9 ms, 16 inversion times (Ti): 35-1500 ms, field of view (FOV): 25.6 mm^2^ (200 μm)^2^, 1 mm thick single slice, acquisition time: 11:44 min). Time Of Flight (TOF) angiography was acquired with a 20 mm diameter quadrature volume coil (RAPID Biomedical, Germany) (TR/TE: 14/2.1 ms, FOV: 20×20×16 mm^3^ (100 μm)^3^, acquisition time: 4:12 min). Spectroscopy, T_2_, susceptibility weighted imaging (SWI), rsfMRI, and dMRI were acquired with a transmit/receive cryocoil (Bruker BioSpin, Germany). A STimulated Echo Acquisition Mode (STEAM) sequence was used for spectroscopy (TR/TE: 2500/3 ms, VAPOR water suppression, 8 mm^3^ striatal voxel), the T_2_ sequence was Rapid Acquisition Relaxation Enhancement (RARE) (TR/TE: 3100/33 ms (100μm^2^), 0.5mm slice thickness), SWI (TR/TE: 700/18 ms, FOV: 75 μm^2^, 0.5 mm slice thickness), a GE-EPI sequence was used for rsfMRI (TR/TE: 1000/10 ms, 300 repetitions, FOV: 19.2×12 mm^2^ (150 μm^2^), 12 contiguous slices @ 0.75 mm) and the dMRI data was acquired with an isotropic EPI-DTI sequence (TR/TE: 7500/18 ms, bvals: 0,1000 s/mm^2^, 126 directions, gradient duration/separation: 2.5/8.1 ms, FOV: 16.2×16.2 mm^2^ (225μm^2^), 58 contiguous slices @ 0.225 mm, acquisition time: 65:30 min).

### Image analysis

CBF maps were calculated via the T_1_ method (Boehm-Sturm et al., 2017), and brain/hippocampal volumes obtained from T_2_ images via manual delineation (ImageJ, NIH, USA). The regions of interest generated from manual segmentation were converted to 3D volumes. Fractal dimensionality was assessed using the calcFD toolbox (Madan and Kensinger, 2017).

All rsfMRI data was converted to NIfTI format using Bru2nii (https://github.com/neurolabusc/Bru2Nii) and bias field corrected using N4BiasFieldCorrection (https://github.com/bigbigbean/N4BiasFieldCorrection) (Tustison et al., 2010). All linear within-subject registrations among structural T_2_ and functional EPI images was performed using affine transforms in Advanced Normalization Tools (ANTs version 2.2.0) (https://github.com/ANTsX/ANTs). ANTs was also used for all non-linear registrations to the Allen Mouse Brain Atlas (https://mouse.brain-map.org/static/atlas). Skull stripping was performed with template brain masks warped into individual subject space. The FMRIB Software library (FSL) (www.fmrib.ox.ac.uk/fsl) was used for rsfMRI data analysis. Correction for image drift and motion was performed with MCFLIRT (Jenkinson et al., 2002). Data was smoothed by the full width at half maximum (0.7mm kernel) and MELODIC was used for independent component analysis (ICA) (Bajic et al., 2017). Neural-driven signal was manually classified at the single subject level as predominantly low frequency, predicted 1.5 – 3 % of the data variance, exhibited no sudden jumps, and activity components needed to be clustered in gray matter (not vasculature, ventricles, or non-brain tissue) (Griffanti et al., 2017). Subsequently, 59 components were produced at the group level, and 19 were manually eliminated as noise according to the same criteria. Dual regression was performed for 14 components of interest followed by voxelwise testing for group differences using FSL’s random permutation testing tool. The p-values in the resulting clusters were Bonferonni corrected (14 comparisons). Masks for seed-based connectivity were generated from the Allen Mouse Brain Atlas or from ICA components.

The MATLAB (version R2019a) toolbox FSLNets (version 0.6) (www.fmrib.ox.ac.uk/fsl) was used to generate all rsfMRI weighted connectivity matrices (Smith et al., 2013). For visualization of connectivity matrices, glass brains were prepared in ParaView (version 5.6.0) (https://www.paraview.org/) from 3D renderings of the surface of the mouse brain prepared from a whole brain mask in ITK-SNAP (version 3.6.0) (http://www.itksnap.org/pmwiki/pmwiki.php) (Avants et al., 2014). Individual components were manually thresholded and exported as binarized masks into ParaView. These masks were then converted into spheres, and z score edges were added and color coded manually. Subsequently, a proportional threshold was applied across the connectivity matrices that ensured the highest small world value (highest clustering and shortest path lengths) was preserved while maintaining matrix density (Fornito et al., 2016; Watts and Strogatz, 1998). The MATLAB (version R2018a) Brain Connectivity Toolbox (https://www.nitrc.org/projects/bct) was used for graph theory and global efficiency, modularity and transitivity were calculated for each animal as well as nodal local efficiency (Rubinov and Sporns, 2010). Circlize visualization in R (https://github.com/jokergoo/circlize) was used to generate the circular views of connectome nodes and edges based on correlation strength (r).

All dMRI data was converted to NIfTI format using Bruker2nifti (https://github.com/SebastianoF/bruker2nifti) (Ferraris et al., 2017) and N4BiasFieldCorrection was employed. B0 template creation and linear affine registration of individual subject data to the B0 template was performed with ANTs, as were all non-liner transforms to the Allen Mouse Brain Atlas. Template brain masks were used for skull stripping via warping into individual subject space. Image pre-processing, correction, and analysis were performed with FMRIB’s Diffusion Toolbox (FDT) (https://fsl.fmrib.ox.ac.uk/fsl/fslwiki/FDT) (Jbabdi et al., 2012). Scalar maps were generated using DTIFIT. Tract

Based Spatial Statistics (TBSS) was used to produce a mean FA map and a white matter skeleton (threshold of 0.4). This was used for voxelwise comparisons of tissue microstructure between groups with Randomise (using the Threshold Free Cluster Enhancement (TFCE) option to define clusters and the family wise error (FWE) corrected p-value option to control for multiple comparisons). For tractography, Bedpostx (Jbabdi et al., 2012) was used to fit a fibre orientation model to each voxel. Probtrackx2 was used with the probabilistic tractography network option (Hernandez-Fernandez et al., 2019). Brain regions were used as seeds and 5000 seeds were placed in each voxel. The step length was half of the voxel size, the angular threshold was 50 degrees, and streamlines were counted if they passed through another atlas region. The resulting structural connectivity matrices reflected the number of streamlines connecting all pairs of regions, and a log_10_ transform was used to normalize for the distance bias in tractography (Ercsey-Ravasz et al., 2013). Graph theory was performed as per the rsfMRI data included appropriate proportional threshold.

Spectroscopic metabolite concentrations were calculated using water scaling to signal in a water-unsuppressed reference scan in LCModel and values with a standard deviation < 25 % were excluded (1 sham and 2 hypoperfused mice at 3 months).

### Statistical analysis

Data is expressed as mean ± standard deviation. The Kolmogorov-Smirnov test established normal distribution (SPSS, IBM, UK). CBF, volumetry, water maze, and novel object recognition data were compared using a mixed two-way repeated measures ANOVA. Graph theory metrics were compared with t-tests or non-parametric equivalents (Mann Whitey or Wilcoxon signed rank) for rsfMRI, or mixed two-way repeated measures ANOVA for dMRI. To correct for multiple comparisons for spectroscopy metabolites and local graph theory parameters, false discovery rate significance thresholds were calculated for all ranked p values and used to correct them (Benjamini and Hochberg, 1995).

## Results and Discussion

### Lacunar infarction, microbleeds and atrophy are observed in mice with vascular cognitive impairment

We employed a well-characterized mouse model of vascular cognitive impairment produced via hypoperfusion to the brain (Coltman et al., 2011; Shibata et al., 2004). We observed additional features of small vessel disease including lacunar infarctions in the striatum in 36 % of hypoperfused mice at 24 hrs, and two animals exhibited thalamic microbleeds by 6 months in susceptibility weighted imaging (SWI) (Fig1A, SFig1A).

**Fig1.**
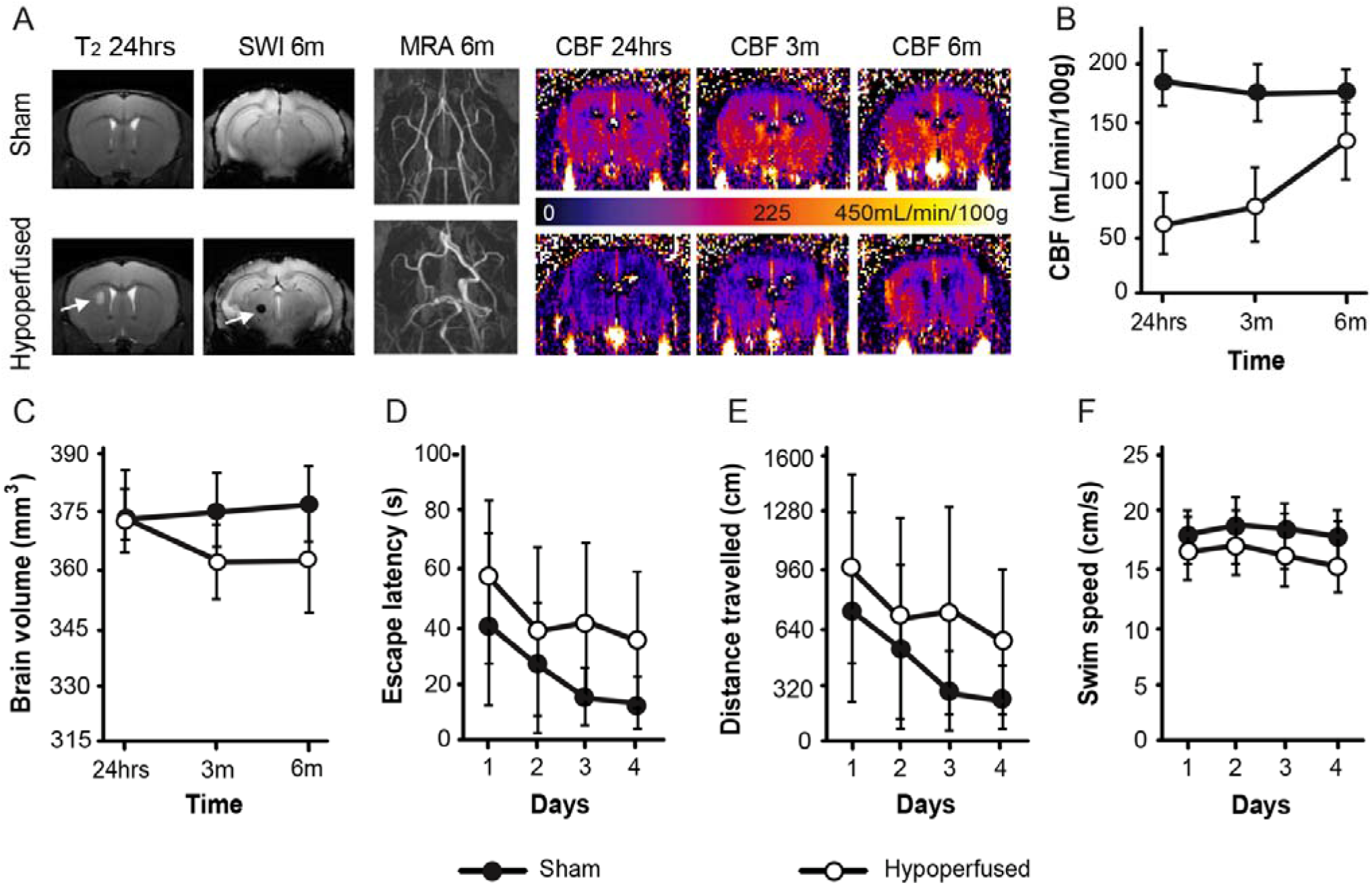
Hypoperfusion causes lacunar infarction, microbleeds and brain atrophy despite recovery of CBF. (A) Representative T_2_ and susceptibility weighted images (SWI) at 24 hrs and 6 months (arrows indicate a lacunar infarction and a microbleed), alongside magnetic resonance angiography (MRA) images of the Circle of Willis at 6 months, and cerebral blood flow (CBF) maps at 24 hrs, 3 and 6 months (scale bar corresponds to CBF). (B) CBF (mean ± SD) is lower in the hypoperfused group (n=6) and these animals exhibited brain atrophy (C). In a water maze cue task at 6 months, hypoperfused mice exhibited longer escape latencies (D), swam greater distances (E), and were slower (F) than sham mice (n=10).

Cerebral blood flow (CBF) remained stable across 6 months in shams. CBF was decreased in hypoperfused mice at 24 hrs, but improved between 3-6 months (time: F(2,26)5.2, p=0.013, group: F(1,13)133.9, p=0.0001, interaction: F(2,26)7.9, p=0.002) (Fig1A-B). This implies that the hypoperfused brain is capable of autoregulation to restore CBF, and has been previously reported by several groups (Füchtemeier et al., 2015; Nishio et al., 2010; Shibata et al., 2004). We observed Circle of Willis remodeling (increased tortuosity and dilation) in all of the hypoperfused mice via magnetic resonance angiography (MRA) (Fig1A), which may underpin the CBF recovery. Despite CBF recovery, the hypoperfused group exhibited significant brain atrophy as measured by a decrease in brain volume (time x group interaction: F(2,26)3.6, p=0.04) (Fig1C). Atrophy was also observed in the hippocampus (Supplementary Results).

### Behavioral deficits in mice with vascular cognitive impairment suggest visual disturbance

A cued task (visible platform) was performed in the water maze at 6 months. While all mice escaped quicker and travelled less distance, faster over the 4 test days (significant main effects of time: F(3,42)13.4, p=0.0001, F(3,42)11.7, p=0.0001, F(3,42)3.4, p=0.027, respectively), the hypoperfused mice took longer to escape, travelled greater distances and swam slightly slower than shams (significant main effects of group: F(1,14)37.6, p=0.0001, F(1,14)33.8, p=0.0001, F(1,14)7.8, p=0.015, respectively) (Fig1D-F). Similar results were observed in a place task (hidden, fixed platform) (SFig2A-C) and hypoperfused mice made fewer returns to the target quadrant during the probe trial (removed platform) (SFig1D-E). This is consistent with a previous report of spatial working and reference memory deficits in the water maze after 6m of hypoperfusion (Holland et al., 2015). Our findings could be interpreted as memory deficits, but as the same effect was observed when the platform is visible, this suggests that the hypoperfused animals may also experience visual impairments. Indeed, retinal dysfunction has been reported in this model that is variable over the course of 6 weeks (Crespo-Garcia et al., 2018; Manouchehrian et al., 2015).

### Characterization of the functional mouse connectome depicts known species-specific hubs

We used a data-driven approach, independent component analysis (ICA), to identify regions with coherent resting state activity (SFig3) (Jonckers et al., 2011). We obtained 59 group-ICA components (n=16); 19 of these were manually classified as noise (Griffanti et al., 2017) (Supplementary Methods). Hierarchical clustering of the 40 remaining components according to correlation strength revealed four clusters (color-coded in Fig2, SFigs4-5).

**Fig2.**
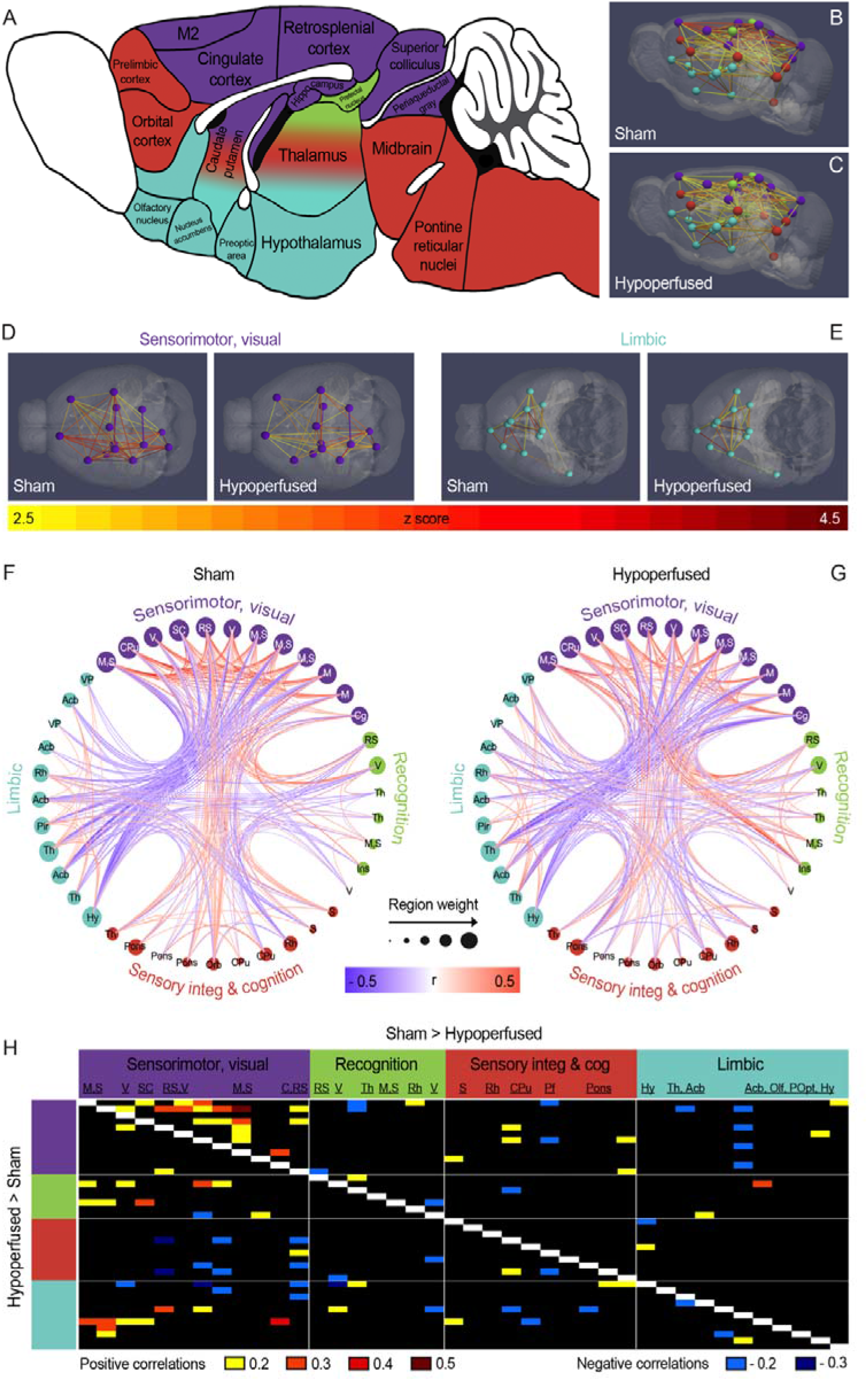
Functional connectivity changes in hypoperfused mice at 6 months. (A) Sagittal mouse brain cartoon (Paxinos and Franklin, 2001) with four color-coded clusters of networks: purple-sensorimotor, visual; green-recognition; red-sensory integration and cognition; and blue-limbic. (B-C) Sagittal 3D and superior 3D views of the sensorimotor, visual and (D) inferior views of the limbic cluster (E) in each group. (F-G) Circular views of connectome nodes and edges displaying connectivity in clusters for each group. Note: nodes represent ICA components and node size reflects the number of edges stemming from each node. (H) Connectivity matrix displaying the strength of the correlation differences between groups (top indicates stronger correlations in shams and bottom stronger correlations in the hypoperfused group). Abbreviations: motor (M), sensory (S), visual (V), retrosplenial (RS), cingulate (C), frontal association (FrA), rhinal (Rh), and prefrontal (Pf) cortices, superior colliculus (SC), thalamus (Th), caudate putamen (CPu), hypothalamus (Hy), nucleus accumbens (Acb), pre-optic nuclei (POpt), olfactory areas (Olf).

The largest and most highly correlated cluster was primarily cortical, and components spanned the sensory, motor, and visual cortices, as well as the retrosplenial and cingulate cortices (Fig2A-D, SFig4: purple). There were also components in the dorsolateral caudate putamen, hippocampus, superior colliculus and peri-aqueductal gray. The sensorimotor, visual cluster was anti-correlated with a ventral cluster that consisted of limbic structures: hypothalamus, nucleus accumbens, ventral pallidum and thalamus, olfactory nucleus, and preoptic area (Fig2A-E, SFig4: blue). Similar components have been grouped into ventral striatal and thalamic or limbic networks previously (Grandjean et al., 2017; Mechling et al., 2014).

There are only a few reports in which hierarchical clustering has been used to examine the functional connectome. The largest and most highly correlated cluster identified by another study also included the sensorimotor, visual and cingulate (referred to as DMN-like) cortices as well as the hippocampus and cerebellum (Zerbi et al., 2015). This is comparable to our findings, apart from the additional cerebellar components. Additional clusters included subcortical, ventral structures such as the caudate, amygdala and ventral pallidum. A recent study also undertook regional portioning that clustered according to nodal strength (Coletta et al., 2020). Strong cortical hierarchy was observed, associative cortical regions acted as hubs where as highly connected nodes that linked hubs were termed sinks. Even without hierarchical analysis, the strong presence of highly correlated cortical components in mice has been well documented using an ICA approach (Jonckers et al., 2014, 2011; Mechling et al., 2014; Nasrallah et al., 2014). A recent multi-center study combined 17 data sets from different laboratories and observed strong cortical representation with ICA (Grandjean et al., 2020). There was also a consistent rostral-caudal DMN spanning the medial prefrontal to retrosplenial cortices when seed based analysis from the cingulate cortex was employed. This had been reported previously (Stafford et al., 2014), occasionally with an anti-correlated somatosensory region (Gozzi and Schwarz, 2016; Liska et al., 2015; Sforazzini et al., 2014). We observed a similar pattern in our data when the cingulate was seeded (SFig6), though the caudal extent of the default mode network was not as extensive.

### The hypoperfused brain exhibits decreased functional connectivity compared to shams

Interestingly, when the data from each group was temporally concatenated and the cingulate seeded, a strong DMN was only observed in the sham animals; very little correlated activity was observed in the hypoperfused mice (SFig6). While this has yet to be reported in a mouse model of vascular cognitive impairment, a decrease in the DMN has been reported in older mice (Egimendia et al., 2019) and in some DMN nodes in a transgenic mouse model of Alzheimer’s disease with human tau repeat domain aggregates (Green et al., 2019).

There were differences in functional connectivity between the sham and hypoperfused groups in some of the clusters (Fig2D-G). Most notably, correlations among components in the sensorimotor, visual cluster were much weaker in hypoperfused animals (Fig2D-G), suggesting the cortical network is more profoundly affected by vascular dementia. Similarly, a decline in functional connectivity has been detected in the sensorimotor cortices of ArcAβ mice (transgenic Alzheimer’s disease) that was apparent as early as 2 months of age, and preceded plaque accumulation by approximately 4 months (Grandjean et al., 2014). We did not observe any differences in the ventral limbic cluster between sham and hypoperfused groups (Fig2D-G). However, when correlation strengths were quantified, there were more positive correlations between the sensorimotor, visual and limbic clusters in the hypoperfused group (Fig2H). As overall, these two clusters were anti-correlated, this suggests that the negative correlation is reduced in hypoperfused animals.

A hypothesis driven approach was also used to select 14 components of interest for follow up. We focused on components in the visual cortex and the hippocampus along with components that corresponded to regions that are highly connected to both structures: thalamus, hypothalamus, and retrosplenial cortex. We also included components in the caudate and orbital frontal cortex, as we were interested in frontal striatal hubs associated with executive function in the mouse. Dual regression of the group-ICA components and voxelwise testing of the spatial maps from the 14 components was used to probe for functional connectivity differences between groups. The hypothesis was rejected in clusters associated with three of the components: hippocampus, one of the visual cortex components and the retrosplenial cortex (SFig7A). The hippocampal and visual cortex components explained 2.34 % and 2.17 % of data variance, respectively. This suggests differences in functional connectivity between these components and brain clusters is different between sham and hypoperfused mice. However, none of the identified clusters associated with the three components remained significant after corrections for multiple comparisons. Nevertheless, this remains of interest, as the behavioral deficits exhibited by hypoperfused mice in the water maze are likely due to hippocampal damage and/or visual impairments.

Despite the observed connectivity differences, the over-arching functional network characteristics (metrics) remained intact in hypoperfused animals. Both groups exhibited typical small-world network properties with short average path lengths and high clustering. There were no differences in global efficiency, modularity, or transitivity between groups (SFig7B-D), suggesting that the fundamental properties of brain network organization were preserved. While global efficiency (average inverse shortest path length for all pairs of nodes) estimates the brain’s overall ability to integrate information, local efficiency is calculated on the basis of each node. Local efficiency was generally lower across most of the components in the hypoperfused group, though more data is required to determine which regions might be most vulnerable, as none of the components survived multiple comparison corrections. Lower regional local efficiency values suggest that there is little resistance to network failure in those regions, and this has been reported in prefrontal and superior temporal regions in patients with subcortical vascular cognitive impairment (Sang et al., 2018; Yu et al., 2015).

### Re-organization of the functional connectome is accompanied by subtle white matter damage without structural connectivity change

Fractional anisotropy (FA) values in the white matter were compared between groups using voxelwise testing via a white matter skeleton. There were clusters of white matter voxels in the sham animals that exhibited higher FA values at 3m post-surgery (SFig8). Most were in the internal capsule with a few smaller clusters scattered throughout the corpus callosum, though they failed to achieve significance. By 6m, there were substantively more clusters that exhibited higher FA values in the sham group. This was particularly clear across much of the corpus callosum, and internal capsule. Two clusters of voxels had significantly higher FA values in shams, though none remained significant after family-wise error rate correction (SFig8). While other groups have reported significant decreases in FA in the white matter of hypoperfused animals around 1m, the results have not always been consistent across structures (Holland et al., 2011; Saggu et al., 2016), and there are also reports of no FA change in hypoperfused animals (Boehm-Sturm et al., 2017; Füchtemeier et al., 2015). It is possible that the focused region of interest based analysis employed in these studies missed key areas, or that the model may not be sufficiently reproducible among research groups. Our voxelwise approach offers the advantage of being able to observe the white matter in it’s entirety, and the increased number of affected clusters by 6m would support the idea that damage is occurring slowly with time. Indeed, more profound decreases in FA in hypoperfused mice have been reported in the corpus callosum, internal capsule and fimbria when examined out to 6m (Holland et al., 2015).

We used probabilistic tractography and there were no overt differences in the resulting tracts between sham and hypoperfused mice (Fig3). Structural connectivity matrices, based on the number of streamlines, were also predictable. The white matter had the largest degree of structural connectivity (more streamlines) with all other regions. Most cortical regions broadly showed high connectivity and the somatosensory area had the highest number of streamlines with other cortical regions. The midbrain was well connected to the thalamus, hypothalamus, motor and retrosplenial cortices, and cerebellum. The medulla was poorly connected with most other regions except the pons and the cerebellum. From all the regions, the caudate had the highest connectivity with the cortex and diencephalon. There were no edge-wise structural connectivity differences between shams and hypoperfused mice, and the same was true when we adapted the Allen Mouse Brain Atlas to focus on only the parent regions (n= 32) (SFig9).

**Fig3.**
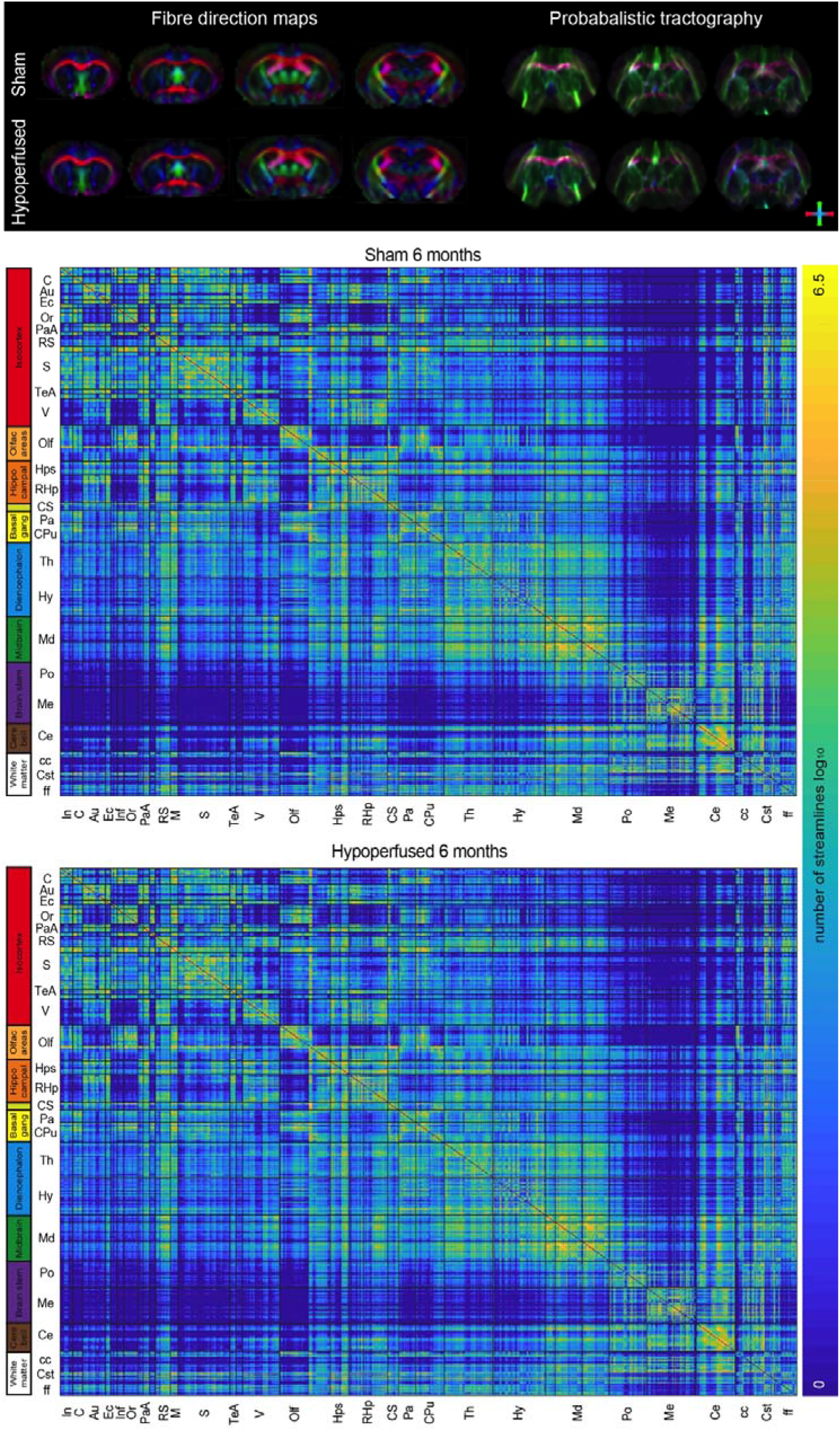
Structural connectivity is intact in hypoperfused mice. Group averaged structural connectivity matrices from 681 seeded brain regions are displayed for sham (top) and hypoperfused (bottom) mice at the 6m timepoint. Note: scale bar corresponds to edge weighting (the log of the number of streamlines between each region). Abbreviations: insular (In), cingulate (C), auditory (Au), ectorhinal (Ec), infralimbic (Inf), orbital (Or), parietal association (PaA), retrosplenial (RS), motor (M), sensory (S), temporal association (TeA), and visual cortices (V), olfactory areas (Olf), hippocampus (Hps), retrohippocampal region (RHp), cortical subplate (CS), ventral pallidum (Pa), caudate putamen (CPu), thalamus (Th), hypothalamus (Hy), midbrain (Md), pons (Po), medulla (Me), cerebellum (Ce), corpus callosum (cc), corticospinal tract (Cst), fornix (ff).

There are only a few papers that examine a dMRI based mouse structural connectome in a disease state. Substantively reduced structural connectivity has been observed in the cuprizone model of demyelination (Saskia Hübner et al., 2017). This is not surprising given the widespread and substantive white matter damage in this model. However, when structural connectivity was compared between APOE3 and APOE4 (a genetic risk factor for Alzheimer’s disease) carrier mice, principle component analysis of pairwise connections revealed interhemispheric sub-networks explained up to 90% of the variation between genotypes (Badea et al., 2019). Interhemispheric connectivity was reduced in the hippocampus and cerebellum of APOE4 when compared to APOE3 mice. While it is possible a similar approach could be used here, our data quality is not directly comparable to the previous studies due our preference for *in vivo* measurements to monitor multiple time-points.

Structural network characteristics were also quantified, and there were no significant differences between sham and hypoperfused mice at either timepoint for global efficiency (SFig10). The utility of global efficiency to predict cognitive decline in patients with small vessel disease may be due to the fact that humans have more white matter and are thus more susceptible when damage is concentrated there. However, considering more chronic timepoints than 6 months may be necessary for widespread inefficiencies in the routes for information exchange to develop across the brain network; this is supported by the voxelwise FA data. It is also possible the rodent brain is capable of compensation that maintains network characteristics. Indeed, no differences in structural or functional global efficiency were observed in five-familial Alzheimer’s disease (5XFAD) mice compared to wildtypes at 23 weeks of age, despite the fact cognitive deficits are observed at this timepoint (Kesler et al., 2018). The 5XFAD mice exhibited higher path lengths, but only at low network densities, and we chose to maintain network density across both groups with a proportional threshold prior to extraction of graph theory parameters as density is well known to influence graph theory parameters. Several nodes exhibited reduced local efficiency in both groups at 6 months (main effect of time) and the corpus callosum, striatum, prelimbic area, hippocampus, corticospinal tract, visual and visual association cortices, pons and hypothalamus all survived multiple correction comparisons. This was accompanied by significant main effects of group in the visual association cortex and hypothalamus, but only the visual association cortex survived. Interestingly, local efficiency was higher in the visual association cortex of the hypoperfused group. This suggests that the visual association cortex is more connected to other brain regions in hypoperfused mice, and may coincide with the decreased functional connectivity observed in the visual cortex.

There are several limitations to the present study including lack of pre-registration, power analysis, and replication. The study was exploratory rather than hypothesis based, but this should not exclude the data from publication.

There is value in a data driven approach, particularly with big data sets as a single hypothesis may be constraining, and interesting findings that could be followed up may be overlooked. We also have a low sample size, and while replication of our findings in a larger cohort would be ideal, we are encouraged that both the imaging and behavioural data suggest similar effects.

## Conclusions

Overall, we observed additional radiological features of small vessel disease in a mouse model of vascular cognitive impairment. Compared to shams, there was little evidence of DMN activity and decreased cortical functional connectivity. The visual cortex and hippocampus appeared to be the most affected nodes, and this suggestion is supported by visual/spatial memory deficits. There was a trend towards subtle white matter damage that became more obvious at the chronic timepoint of 6 months, though overall structural network integrity remained intact.

## Abbreviations

5XFAD: five-familial Alzheimer’s disease
Acb: nucleus accumbens
Au: auditory cortex
C: cingulate cortex
CBF: cerebral blood flow
cc: corpus callosum
Ce: cerebellum
Cst: corticospinal tract
CuP: caudate putamen
CS: cortical subplate
DMN: default mode network
dMRI: diffusion magnetic resonance imaging
Ec: ectorhinal cortex
FA: fractional anisotropy
ff: fornix
FrA: frontal association cortex
Hps: hippocampus
Hy: hypothalamus
ICA: independent component analysis
In: insular cortex
Inf: infralimbic cortex
M: motor cortex
Md: midbrain
Me: medulla
MRA: magnetic resonance angiography
Olf: olfactory areas
Or: orbital cortex
Pa: ventral pallidum
PaA: parietal association cortex
Pf: prefrontal cortex
Po: Pons
POpt: preoptic nuclei
Rh: rhinal cortex
RHp: retrohippocampal region
RS: retrosplenial cortex
rsfMRI: resting state functional magnetic resonance imaging
S: sensory cortex
SC: superior colliculus
SWI: susceptibility weighted images
TeA: temporal association cortex
Th: thalamus
V: visual cortex

