## Supplementary file for "Vascular cognitive impairment in the mouse reshapes visual, spatial network functional connectivity"

**Supplementary Results**

**Lacunar infarction, microbleeds and atrophy are observed in mice with vascular cognitive impairment**

Approximately 36 % of hypoperfused mice exhibited lacunar infarctions at 24 hrs (Fig1A, SFig1A) and the hypoperfused group exhibited significant brain atrophy (Fig1C). Atrophy was also observed in the hippocampus of hypoperfused mice (group: F(1,13)5.4, p=0.037, time: F(2,26)1.7, p=0.208, interaction: F(2,26)1.1, p=0.4). Fractal dimensionality is a measure of structural complexity, and has been shown to be more sensitive to inter-individual differences than volume measurements. When fractal dimensionality was investigated, there were no significant changes over time in the brain (F(1,14)0.6, p=0.457, Greenhouse-Geisser) or the hippocampus (F(1,14)0.6, p=0.451, Greenhouse-Geisser) (SFig1B-C). There were also no significant differences in fractal dimensionality between groups in the brain (group: F(1,14)0.6, p=0.458, interaction: F(1,14)0.584, p=0.458) or the hippocampus (group: F(1,14)0.7, p=0.404, interaction: F(1,14)0.5, p=0.486) (SFig1B-C). The lack of differences in structural complexity of the brain or the hippocampus indicates that, unlike in humans, mice do not have sufficient shape detail for fractal dimensionality.

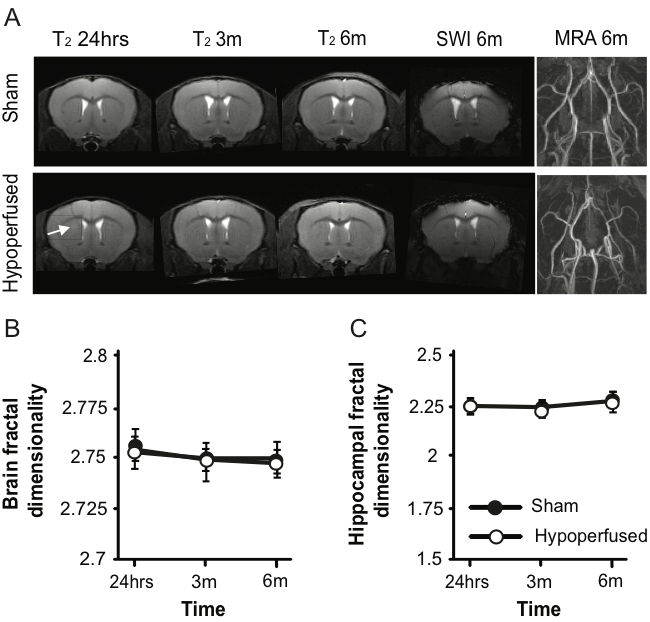

SFig1. Hypoperfusion causes lacunar infarctions and brain and hippocampal atrophy without affecting fractal dimensionality. (A) Representative T_2_ images from a sham and hypoperfused animal (arrow indicates a lacunar infarction) throughout the course of the experiment alongside a susceptibility weighted image (SWI) and a magnetic resonance angiography (MRA) image of the Circle of Willis at 6m. (B-C) There were no significant differences in fractal dimensionality between groups throughout the course of the experiments.

**Behavioral deficits in mice with vascular cognitive impairment suggest visual disturbance**

In a water maze place task (hidden, fixed platform), escape latencies (F(6,84)2.7, p=0.019), total distance travelled (F(6,84)3.9, p=0.002), and swim speed (F(2.9,40.2)2.3, p=0.0001) decreased with time (SFig2A-C). Hypoperfused mice did not escape as quickly (F(1,14)42.8, p=0.0001, no interaction), they travelled greater distances to find the platform (F(1,14)25.6, p=0.0001, no interaction), and swam slower (F(1,14)15.6, p=0.001, no interaction) (SFig2A-C). During the probe trial (removed platform), there were no significant group differences for total time spent in the target quadrant (t(14)1.5, p=0.15) (SFig2D), however, shams swam further (t(14)3.2, p=0.006), faster (t(14)3.2, p=0.006), and made more returns to the platform location (t(14)3.6, p=0.003) (SFig2E). During NOR, all mice explored both objects equally during the first trial, and spent more time with the novel object during the second trial (main effect of time: F(1,14)133.4, p=0.0001).This was not influenced by hypoperfusion (no main effect of group or interaction) (SFig2F). Consistent with previous reports (Patel et al., 2017; Toyama et al., 2014), hypoperfused mice did not exhibit deficits in short term recognition memory in the novel object recognition task (SFig1F).

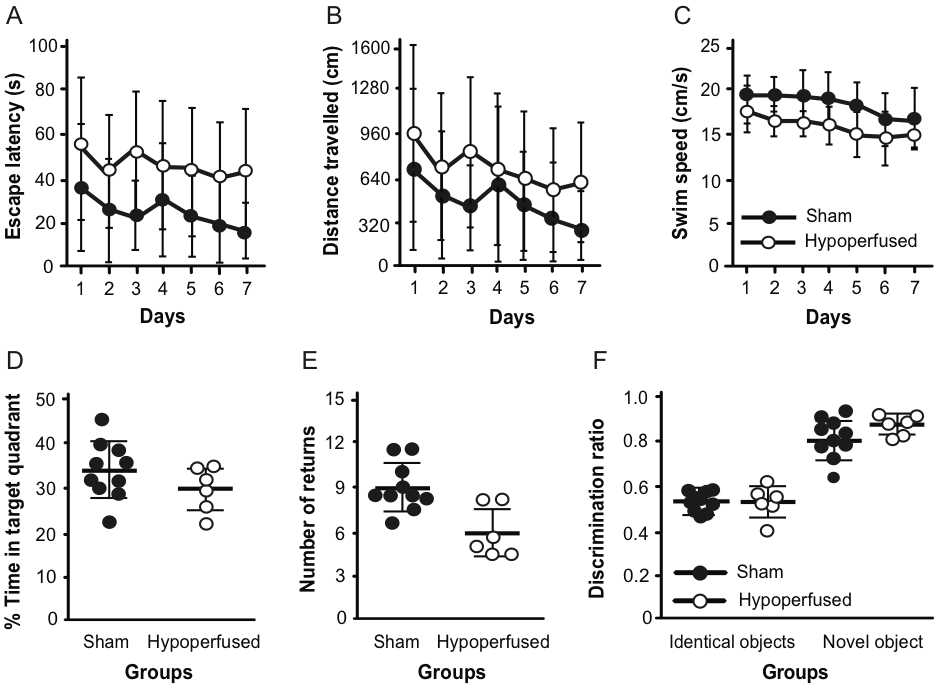

SFig2. Hypoperfused mice exhibit deficits in the water maze without impairments in recognition memory. Escape latency (A), total distance travelled (B) and swim speed (C) in a place task in the Morris water maze. Time spent in (D) and number of returns to the target quadrant (E) during the probe trial with no platform. (F) Discrimination ratio of both groups in the NOR task.

**Characterization of the functional mouse connectome depicts known species-specific hubs**

There is uncertainty surrounding the appropriate number of components for ICA. Specification of low numbers of components (15 and 30) (Jonckers et al., 2011) resulted in large areas of activation with limited regional specificity (SFig3). Ultimately, a data driven approach was selected; principle component analysis chooses the number of components that reflect a predictable amount of variance that can be explained according to data quality.

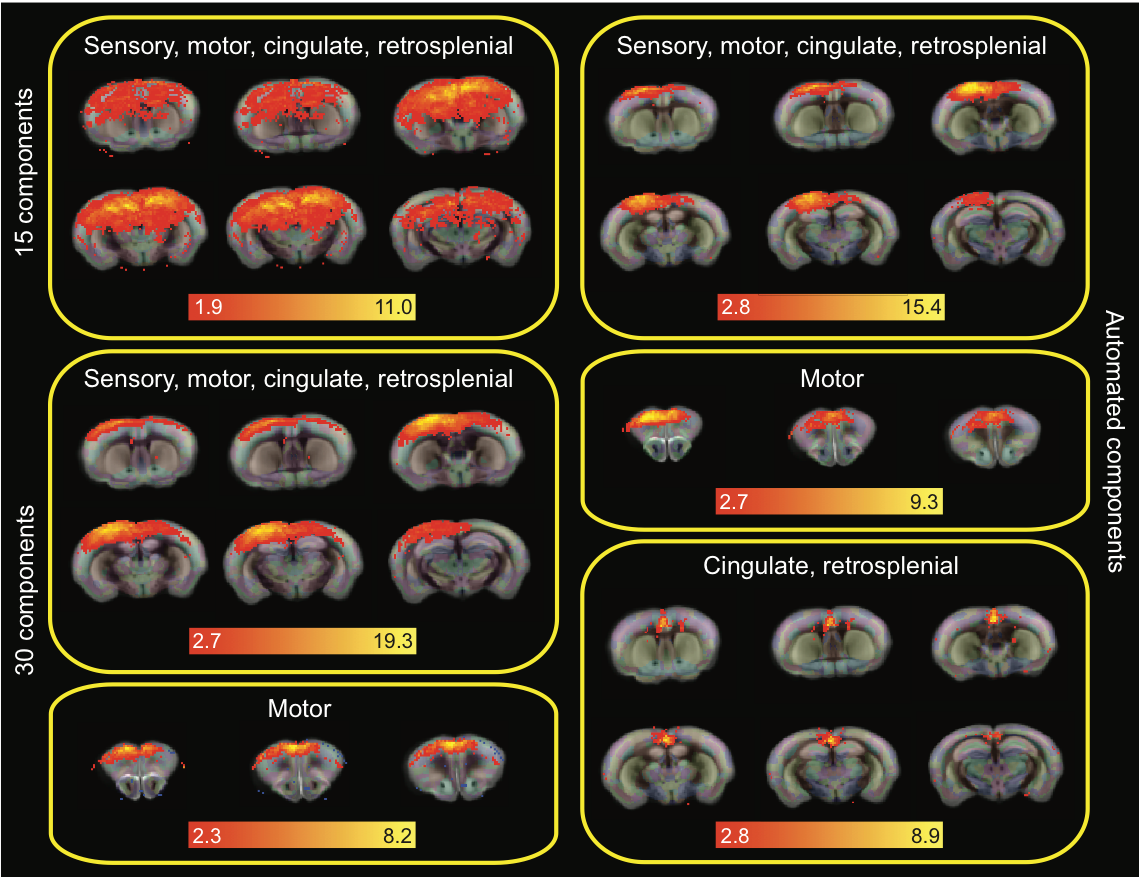

SFig3. Variation in the number of selected components. The same sub-network (sensory, motor, cingulate, and retrosplenial cortices) is depicted under an analysis that specified 15 (top left), 30 (middle and bottom left), and an automatically determined number of components (right). A motor cluster is depicted from the 30 and automated analysis (bottom left and middle right, respectively) alongside additional components in the cingulate and retrosplenial cortices that were detected from the automated analysis. Note: Components are overlaid onto the Allen Mouse Brain Atlas (<http://mouse.brain-map.org/static/atlas>) and scale bars correspond to the z-score.

Hierarchical clustering produced four clusters (Fig2). The sensorimotor, visual cluster was composed of components in the sensory, motor, and visual cortices as well as in the cingulate and retrosplenial cortices (SFig4: purple). There were also sub-cortical components linked to the sensorimotor (dorsolateral caudate and peri-aqueductal gray) and visual systems (superior colliculus, hippocampus). This cluster was anti-correlated with a ventral cluster of limbic components: hypothalamus, nucleus accumbens, ventral pallidum, thalamus, olfactory nucleus, and preoptic area (SFig4: blue). These regions are well known to play a role in homeostatic and social functions, as well as motivation, reviewed in (Castro et al., 2015; Smith et al., 2009; Sokolowski and Corbin, 2012).

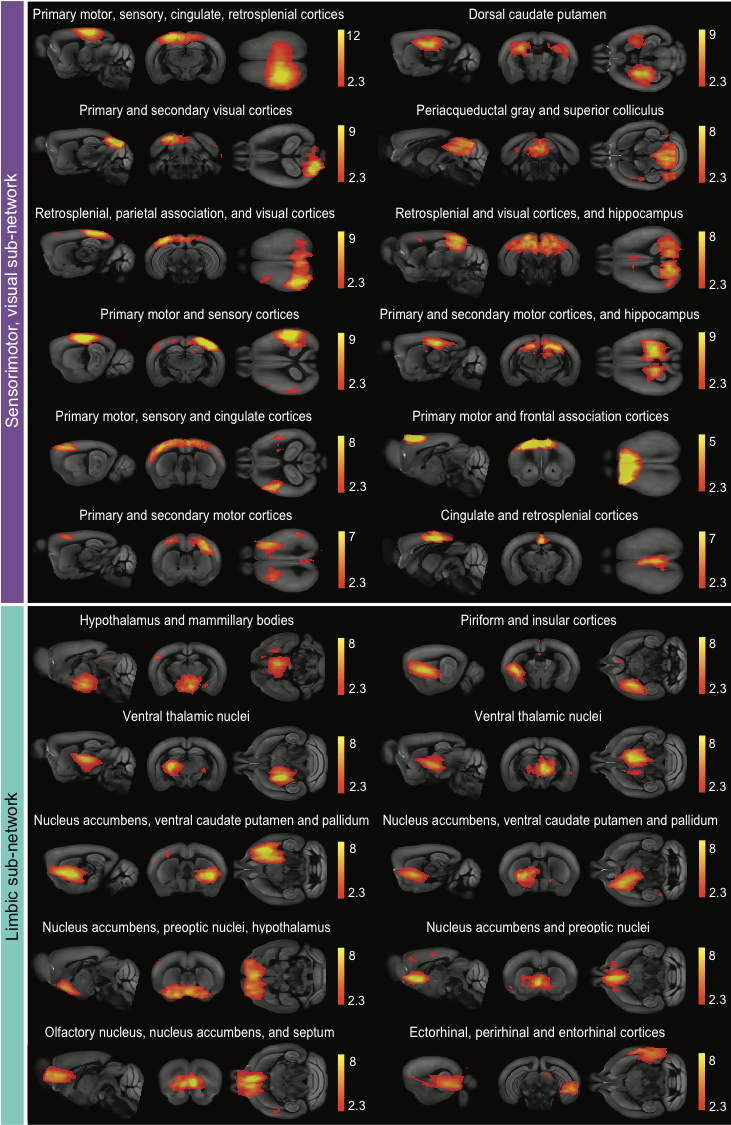

SFig4. The sensorimotor, visual and limbic sub-networks. The sensorimotor, visual (purple) contains 12 and the limbic (blue) sub-network contains 11 independent activity components that are highly correlated (10 pictured). These sub-networks were negatively correlated with each other. Note: Components are overlaid onto the Allen Mouse Brain Atlas (<http://mouse.brain-map.org/static/atlas>) and scale bars correspond to the z-score.

The recognition, and sensory integration and cognition clusters are depicted in SFig5.

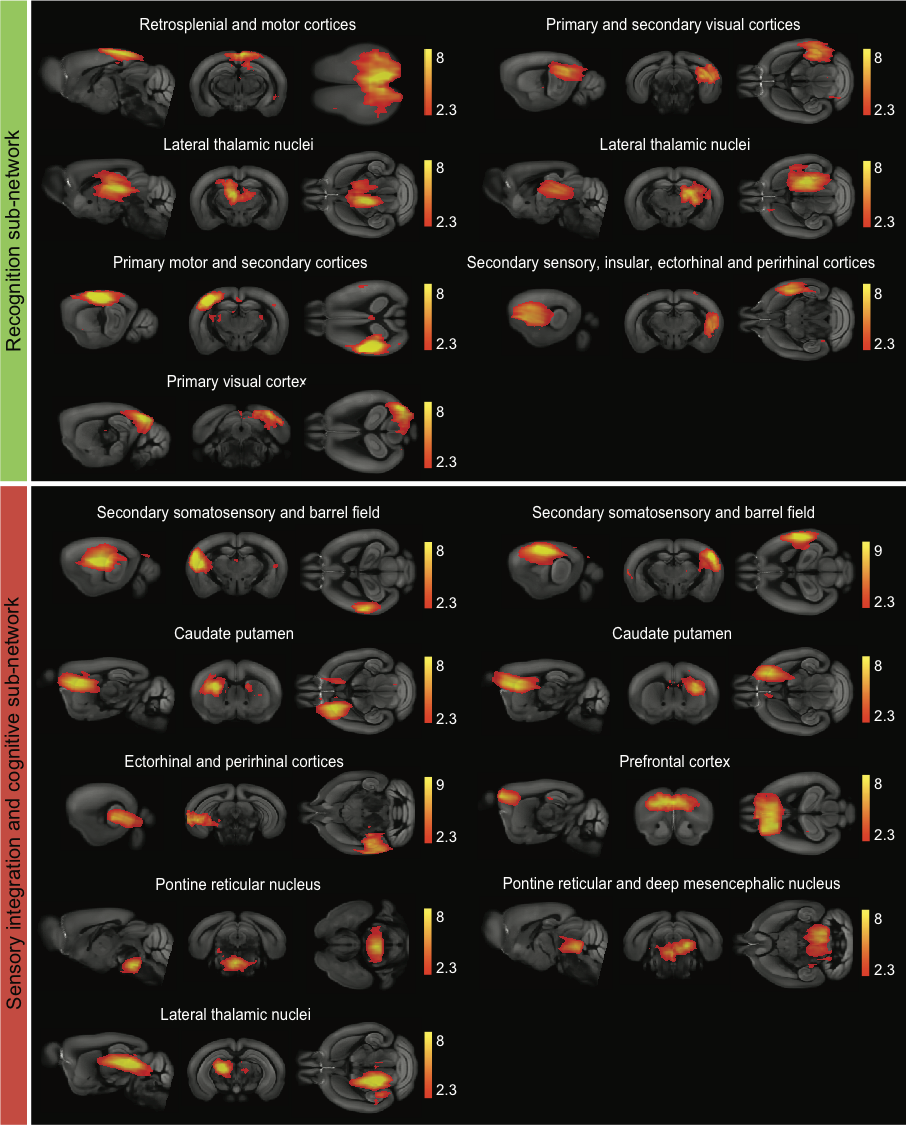

SFig5. The recognition and sensory integration and cognition sub-networks. The recognition (green) contains 7 and the sensory integration and cognition (red) sub-network contains 9 independent activity components that are highly correlated. Note: Components are overlaid onto the Allen Mouse Brain Atlas (<http://mouse.brain-map.org/static/atlas>) and scale bars correspond to the z-score.

The recognition cluster included cortical (retrosplenial, visual, and sensorimotor) and lateral thalamic components (SFig5: green). The retrosplenial cortex has reciprocal connections with the sensory, motor and visual cortices, as well as the anterolateral thalamus. It has been proposed to play a role in integration of information, particularly in the context of spatial navigation (learning locations, recognizing landmarks), reviewed in (Mitchell et al., 2018; Seabrook et al., 2017). There are also reciprocal connections between the visual cortices and lateral thalamic nuclei (Allen et al., 2016).

The sensory association and cognition cluster (SFig5: red) contains activity components in the secondary somatosensory and prefrontal cortices (including the anterior cingulate), as well as the caudate and aspects of the midbrain and pons. The secondary somatosensory cortex is reciprocally connected to the prefrontal cortex. The former is generally considered to be a major integrative structure when it comes to processing sensory information, while the latter is involved with performance adjustment and modulating behaviors, reviewed in (Laubach et al., 2018). There are also projections to the medial caudate from the prefrontal cortex that are associated with sensory and cognitive processes, reviewed in (Voorn et al., 2004). The functions of the pontine tegmental area and deep mesencephalic nucleus are not well characterized, but in addition to being associated with sleep and consciousness, they receive striatal input and may participate in perception, vigilance and awareness associated with cognitive basal ganglia behaviors (Rodriand́guez et al., 2001).

**The hypoperfused brain exhibits decreased functional connectivity compared to shams**

Quantification of the strength of the correlations between groups among the clusters revealed interesting patterns (Fig2H) that were also visible in the circle visualizations of the connectomes (Fig2D,E). The hypoperfused group exhibited more positive correlations between the sensorimotor, visual and recognition clusters, and more negative correlations between the sensorimotor, visual and sensory integration and cognition clusters. This may represent some form of compensation for the overall decline in connectivity in the sensorimotor, visual cluster. We were particularly interested in a small hub consisting of the prefrontal cortex and the caudate putamen, as these structures are known to play a role in executive function in rodents. We used a hypothesis driven approach to probe this hub with voxelwise testing, but there were no significant differences between groups.

Several groups have reported a midline rostral-caudal cortical activity pattern that emerges when the cingulate cortex is seeded; this has been referred to as the rodent default mode network (DMN) (Gozzi and Schwarz, 2016; Grandjean et al., 2019; Jonckers et al., 2011; Liska et al., 2015; Nasrallah et al., 2014; Sforazzini et al., 2014). We observed a similar pattern, though activity was slightly more rostral and ventral. However, when the sham and hypoperfused groups were temporally concatenated, there was virtually no DMN activity in the hypoperfused group (SFig6).

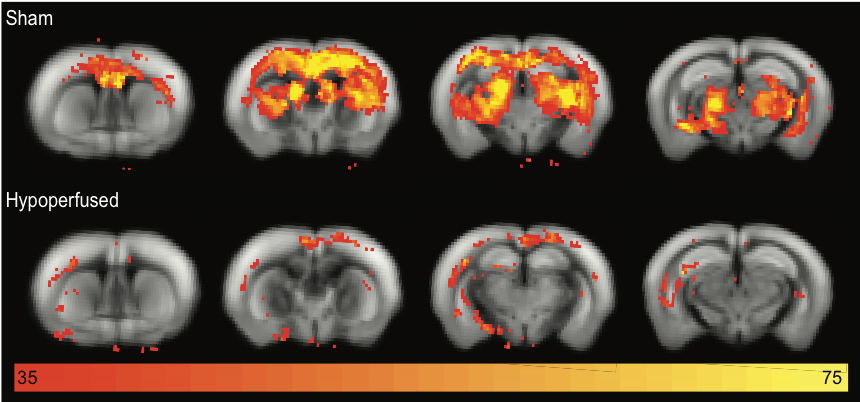

SFig6. The default mode network is reduced in hypoperfused mice. Cingulate seeding produced a strong, primarily cortical, rostral-caudal midline activity pattern (rodent default mode network) in the sham animals that was markedly reduced in the hypoperfused group. Note: Components are overlaid onto the Allen Mouse Brain Atlas (<http://mouse.brain-map.org/static/atlas>) and scale bars correspond to the z-score.

A hypothesis driven approach was used to select 14 components of interest based on regional location (hippocampus, visual and retrosplenial cortices, thalamus, hypothalamus, caudate and orbital frontal cortex). Following dual regression of group-ICA components and voxelwise testing on the spatial maps of the 14 selected components, three of the components had clusters of voxels in which the hypothesis was rejected: the hippocampus, the retrosplenial cortex and one of the visual cortex components (SFig7A). This suggests differences in functional connectivity between these components and brain clusters, between sham and hypoperfused mice, despite a lack of significance after 14 multiple comparison corrections.

When overall network characteristics were estimated from the rsfMRI data, there were no significant differences between groups for global efficiency (U=26, p=0.713), modularity (the breaking down of the network into groups of nodes based on maximum within group and minimal between group connections: t(14)0.2, p=0.854), or transitivity (U=16, p=0.147) (SFig7B-D).

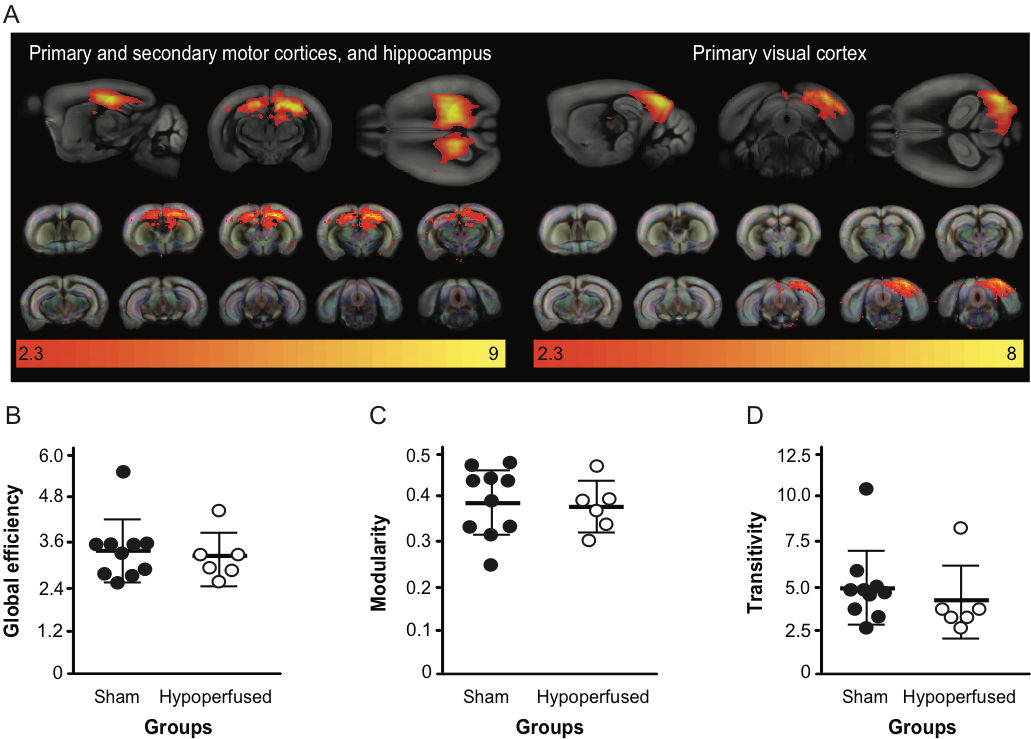

SFig7. The hippocampus and visual cortex were the most affected brain regions though the entire network exhibited preserved characteristics. (A) Hippocampus and visual cortex components exhibited connectivity differences between the sham and hypoperfused groups. Note: Components are overlaid onto the Allen Mouse Brain Atlas (<http://mouse.brain-map.org/static/atlas>) and scale bars correspond to the z-score. (B) Global efficiency, (C) modularity and (D) transitivity network properties in sham and hypoperfused groups.

**Re-organization of the functional connectome is accompanied by subtle white matter damage without structural connectivity change**

A voxelwise approach was applied to the dMRI data to compare FA values across the white matter between groups. FA values were higher in clusters in the internal capsule and corpus callosum in sham animals at 3m (SFig8), though none of the voxels achieved significance. By 6m, shams higher FA values in most of the corpus callosum and internal capsule. Two clusters of voxels were significantly higher in shams, though this was not the case after family-wise error rate correction (SFig8). Overall, this suggests white matter damage is an evolving process in this model.

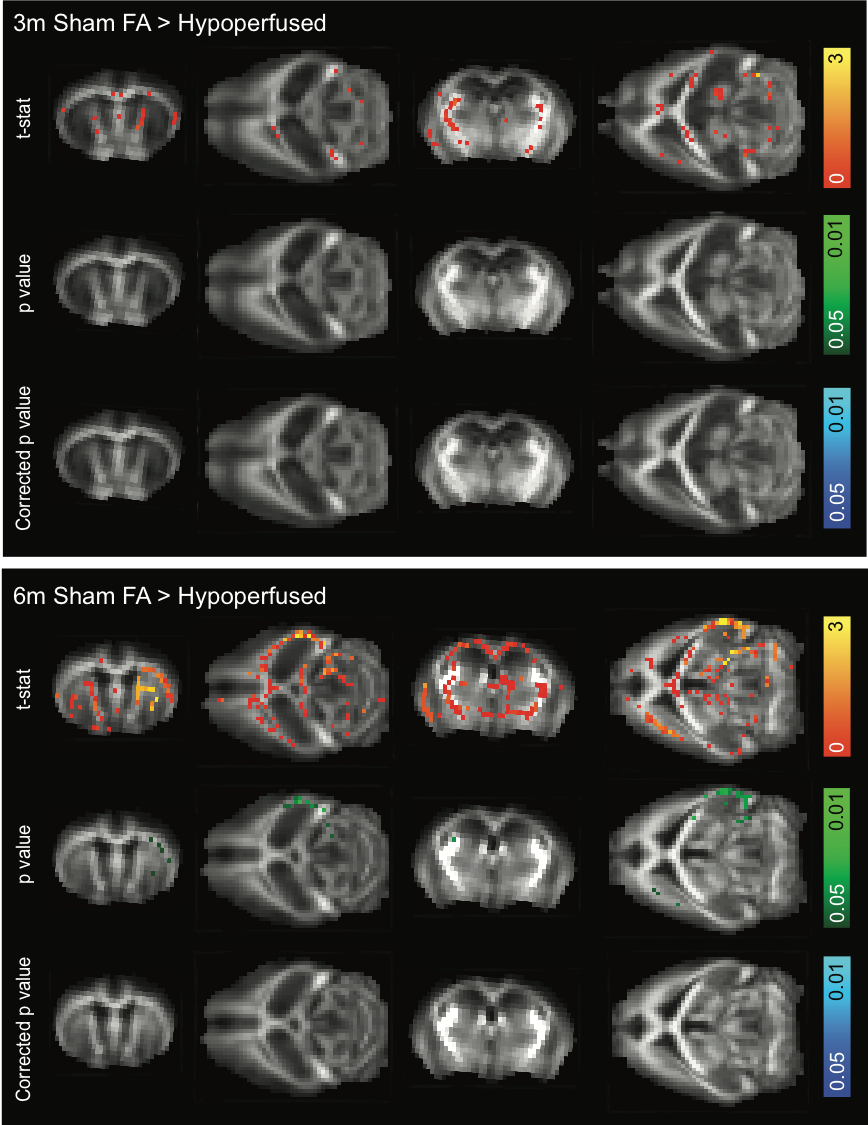

SFig8. White matter is only subtly affected by hypoperfusion. Sham animals exhibited higher FA values in white matter voxels at 3 and especially 6m. Note: orange scale bar represents the t-statistic, green scale bar represents the p value, and blue scale bar represents the corrected p-value overlaid onto the mean FA map from all mice at each timepoint (none of the p values survived multiple comparisons).

Probabilistic tractography was performed, and there were no overt differences the resulting tracts or structural connectivity matrices between sham and hypoperfused mice (Fig3). However, the 681/1206 Allen Mouse Brain Atlas regions that survived down-sampling are still well beyond the resolution of the dMRI. A previous report directly compared a connectome generated from axonal tracing experiments (the Allen Mouse Brain Connectivity Atlas) (Oh et al., 2014) to a dMRI based mouse structural connectome (Chen et al., 2015). In addition to establishing ideal tractography parameters, there was a 90% similarity between the two structural connectomes when a courser atlas parcellation of 96 regions was used. Another report compared the same neural tracing atlas to a probabilistic tractography based structural connectome generated using the same methods as the current study (Calabrese et al., 2015). Structural connectivity was almost indistinguishable (99% correlation) when atlas regions were collapsed into only the parent structures. Therefore, we combined the Allen Mouse Brain Atlas regions into only the parent structures (32 regions), though there were still no connectivity differences observed between groups at either 3 or 6 months (SFig9).

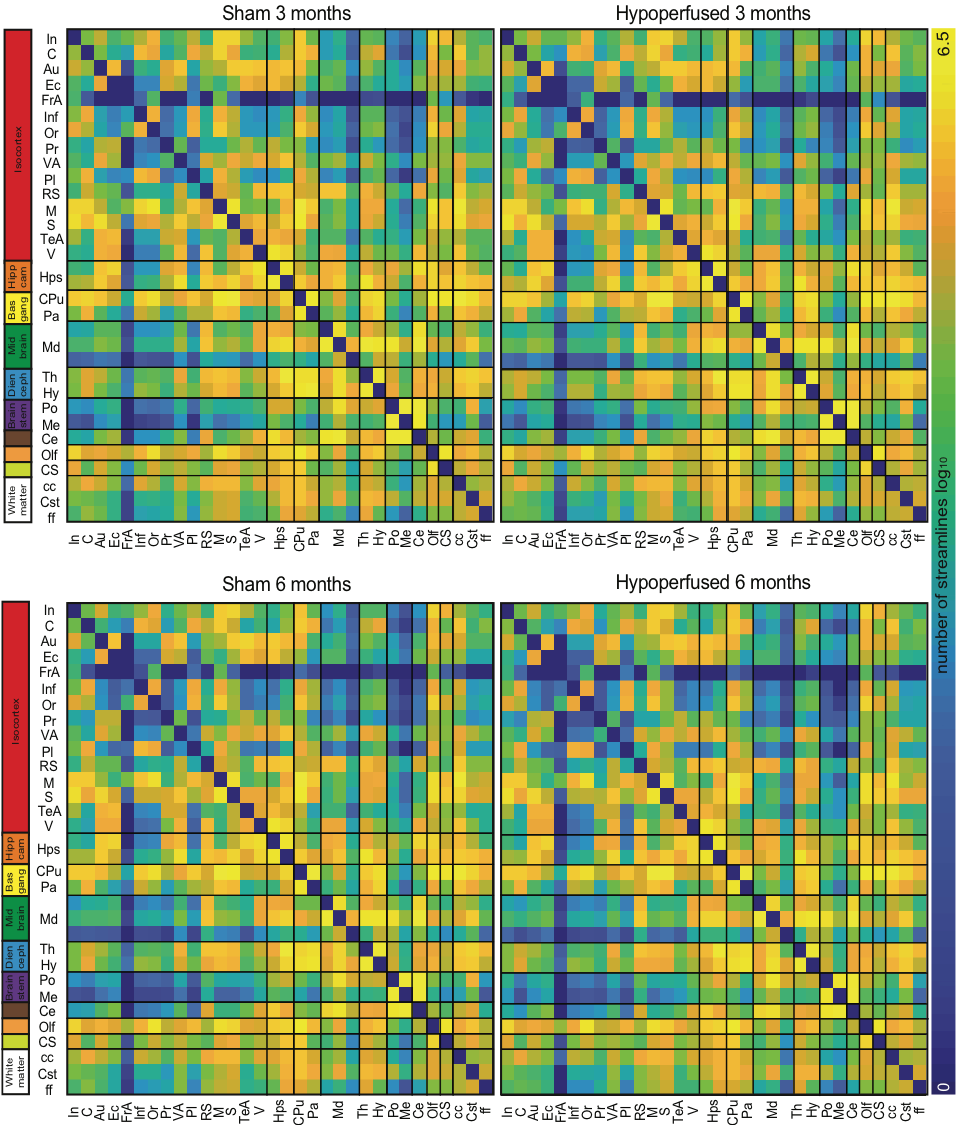

SFig9. Structural connectivity is intact in hypoperfused mice at both timepoints. Structural connectivity matrices from 32 seeded parent regions are displayed for sham and hypoperfused mice at 3 and 6m. Note: scale bar corresponds to edge weighting (the log of the number of streamlines between each region). Summary regions (left) correspond to higher level collective groups from the Allen Mouse Brain Atlas (<http://mouse.brain-map.org/static/atlas>) (Isocortex, Hippocampal formation, Cerebral nuclei (basal ganglia), Midbrain, Hindbrain (diencephalon and brain stem), Cerebellum, Olfactory areas, Cortical subplate, and Fibre tracts (white matter). Abbreviations: insular (In), cingulate (C), auditory (Au), ectorhinal (Ec), frontal association (FrA), infralimbic (Inf), orbital (Or), perirhinal (Pr), visual association (VA), prelimbic (Pl), retrosplenial (RS), motor (M), sensory (S), temporal association (TeA), and visual (V) cortices, hippocampus (Hps), caudate putamen (CPu), ventral pallidum (Pa), midbrain (Md), thalamus (Th), hypothalamus (Hy), pons (Po), medulla (Me), cerebellum (Ce), olfactory areas (Olf), cortical subplate (CS), corpus callosum (cc), corticospinal tract (Cst), and fornix (ff).

Graph theory was used to examine overall structural network characteristics. There were no significant differences in global efficiency (group: F(1,14)0.61, p=0.447, time: F(1,14)0.25, p=0.625, time x group interaction: F(1,14)0.025, p=0.876), or modularity (group: F(1,14)2.6, p=0.127, time: F(1,14)2.9, p=0.109, time x group interaction: F(1,14)0.02, p=0.890). (SFig10A-B). While there was no main effect of group for transitivity (group: F(1,14)0.79, p=0.388), this parameter decreased in both groups at 6m (time: F(1,14)20.4, p=0.0001, time x group interaction: F(1,14)0.007, p=0.936) (SFig10A-C). Transitivity reflects the extent to which a connection can be observed between 2 nodes that each share a connection with another neighboring node. This feature is typically high in networks, and a decrease with age may imply a reduced ability to compensate for loss of function due to fewer alternative pathways to share information.

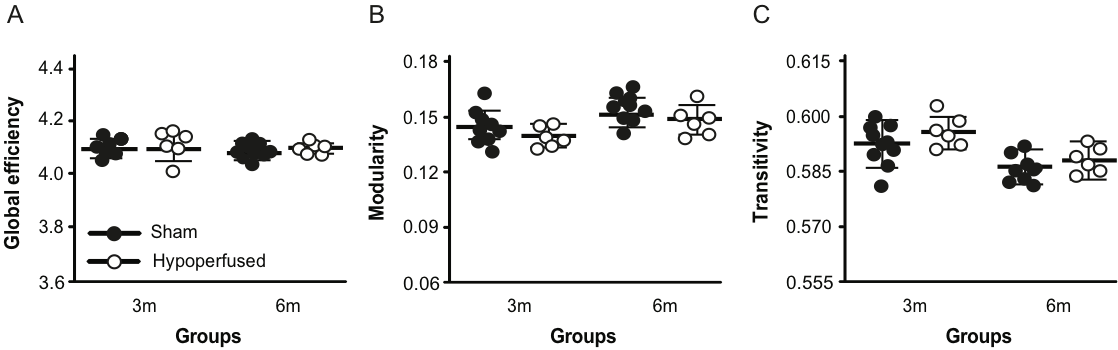

SFig10. Hypoperfused mice exhibit intact structural network characteristics. (A)

(B) Global efficiency, (B) modularity and (C) transitivity network properties in sham and hypoperfused groups at 3 and 6m.

**Spectroscopy revealed limited evidence for metabolic changes in hypoperfused mice**

Metabolite concentrations were measured using localized MR spectroscopy in the striatum. Interestingly, while there were no significant differences in metabolite between the sham and hypoperfused groups, glycerophosphocholine and glutamate and glutamine increased in both groups with age and alanine and lactate decreased (STable1). Taurine exhibited a significant time by group interaction in that it increased in the sham group and decreased in the hypoperfused group with time.

| Table 1. Brain metabolite concentrations (mM) in sham and hypoperfused mice across 6 months. | | | | | | | | | |
| --- | --- | --- | --- | --- | --- | --- | --- | --- | --- |
|  | S ham n(%) | | |  | Hypoperfused n(%) | | |  |  |
|  | 3months  7(70) | 6months  9(90) | %  change |  | 3months  4(36) | 6months  6(55) | %  change |  | p |
|  |  |  |  |  |  |  |  |  | value |
| N-acetylaspartate (NAA) | 2.7 ± 0.09 | 2.78 ± 0.14 | 4.3 |  | 2.74 ± 0.28 | 2.72 ± 0.09 | -0.8 |  |  |
| Creatine (Cr) | 1.76 ± 0.09 | 1.69 ± 0.15 | -4.2 |  | 1.58 ± 0.24 | 1.70 ± 0.11 | 7.5 |  |  |
| Phosphocreatine (PCr) | 1.15 ± 0.15 | 1.36 ± 0.14 | 16.3 |  | 1.29 ± 0.32 | 1.33 ± 0.13 | 2.7 |  |  |
| Glycerophosphocholine (GPC) | 0.22 ± 0.04 | 0.31 ± 0.07 | 1.7 |  | 0.24 ± 0.07 | 0.31 ± 0.05 | 26.4 |  | F(1,9) = 8.372, p = 0.018 * |
| Phosphocholine (PCh) | 0.48 ± 0.05 | 0.44 ± 0.07 | -7.4 |  | 0.47 ± 0.12 | 0.41 ± 0.09 | -12.2 |  |  |
| Glutamate (Glu) and Glutamine (Gln) | 4.11 ± 0.17 | 4.83 ± 0.26 | 16.1 |  | 4.20 ± 0.35 | 4.75 ± 0.19 | 12.3 |  | F(1,9) = 20.76, p = 0.001* |
| Inositol (Ins) | 1.99 ± 0.18 | 1.91 ± 0.19 | -4.0 |  | 2.15 ± 0.32 | 1.99 ± 0.11 | -7.5 |  |  |
| Glucose (Glc) | 0.68 ± 0.32 | 0.82 ± 0.19 | 17.8 |  | 0.60 ± 0.26 | 0.76 ± 0.12 | 23.6 |  |  |
| N-Acetylaspartylglutamate (NAAG) | 0.02 ± 0.02 | 0.04 ± 0.04 | 61.3 |  | 0.02 ± 0.03 | 0.02 ± 0.03 | -14.4 |  |  |
| Taurine (Tau) | 4.44 ± 0.16 | 4.74 ± 0.33 | 6.4 |  | 4.55 ± 0.28 | 4.43 ± 0.16 | -2.6 |  | F(1,9) = 6.568, p = 0.030 ‡ |
| Alanine (Ala) | 0.76 ± 0.12 | 0.56 ± 0.10 | -30.7 |  | 0.74 ± 0.17 | 0.54 ± 0.06 | -32.1 |  | F(1,9) = 12.52, p = 0.006 * |
| Lactate (Lac) | 2.38 ± 0.49 | 1.86 ± 0.28 | -24.4 |  | 2.12 ± 0.39 | 1.66 ± 0.16 | -24.1 |  | F(1,9) = 7.977, p = 0.020 * |
| * Indicates significant effect of time, ‡ indicates significant time x group interaction | | | | | | |  |  |  |
